# *In Vitro* Activity of Itraconazole Against SARS-CoV-2

**DOI:** 10.1101/2020.11.13.381194

**Authors:** Ellen Van Damme, Sandra De Meyer, Denisa Bojkova, Sandra Ciesek, Jindrich Cinatl, Steven De Jonghe, Dirk Jochmans, Pieter Leyssen, Christophe Buyck, Johan Neyts, Marnix Van Loock

**Author notes:** Corresponding Author, **Address for Correspondence** Marnix Van Loock, PhD, Infectious Diseases & Vaccines Discovery, Janssen Pharmaceutica NV, Turnhoutseweg 30, 2340 Beerse, Belgium.

## Abstract

**Background:** As long as there is no vaccine available, having access to inhibitors of SARS-CoV-2 will be of utmost importance. Antivirals against coronaviruses do not exist, hence global drug re-purposing efforts have been carried out to identify agents that may provide clinical benefit to patients with COVID-19. Itraconazole, an antifungal agent, has been reported to have potential activity against animal coronaviruses.

**Methods:** Using cell-based phenotypic assays, the *in vitro* antiviral activity of itraconazole and 17-OH itraconazole was assessed against clinical isolates from a German and Belgian patient infected with SARS-CoV-2.

**Results:** Itraconazole demonstrated antiviral activity in human Caco-2 cells (EC_50_ = 2.3 μM; MTT assay). Similarly, its primary metabolite, 17-OH itraconazole, showed inhibition of SARS-CoV-2 activity (EC_50_ = 3.6 μM). Remdesivir inhibited viral replication with an EC_50_ = 0.4 μM. Itraconazole and 17-OH itraconazole resulted in a viral yield reduction *in vitro* of approximately 2-log_10_ and approximately 1-log_10_, as measured in both Caco-2 cells and VeroE6-eGFP cells, respectively. The viral yield reduction brought about by remdesivir or GS-441524 (parent nucleoside of the antiviral prodrug remdesivir; positive control) was more pronounced, with an approximately 3 log_10_ drop and >4 log_10_ drop in Caco-2 cells and VeroE6-eGFP cells, respectively.

**Discussion:** Itraconazole and 17-OH itraconazole exert *in vitro* low micromolar activity against SARS-CoV-2. Despite the *in vitro* antiviral activity, itraconazole did not result in a beneficial effect in hospitalized COVID-19 patients in a clinical study (EudraCT Number: 2020-001243-15).

**Highlights:**

- Itraconazole exerted *in vitro* low micromolar activity against SARS-CoV-2 (EC_50_ = 2.3 μM)
- Remdesivir demonstrated potent antiviral activity, confirming validity of the assay
- Itraconazole has since shown no efficacy in a clinical study in hospitalized COVID-19 patients

## Introduction

The rapid spread of the severe acute respiratory syndrome-associated coronavirus-2 (SARS-CoV-2) disease (COVID-19) has been declared a global pandemic.^1,2^ SARS-CoV-2 is a beta-coronavirus, which are enveloped viruses containing single-strand, positive-sense RNA.^3^ Since COVID-19 emerged in humans in late December 2019,^4^ aside from a significant economic loss, more than a million death have been reported worldwide, driving urgently the need to identify potential vaccines and therapies.

Given that therapeutic options for antiviral treatment of SARS-CoV-2 remain limited, research has initially focused on re-purposing available drugs that have demonstrated antiviral activity against coronaviruses. As part of such an initiative, itraconazole was identified as a potential candidate. Itraconazole has been described previously to have activity in an *in vitro* screen using a luciferase reporter-expressing recombinant murine betacoronavirus,^5^ as well as against a feline alphacoronavirus that causes feline infectious peritonitis (FIP).^6^ Furthermore, *in vitro* and *in vivo* activity has been described against other respiratory viruses, such as influenza A and human rhinovirus.^7,8^

Itraconazole is a member of the triazole group of broad-spectrum antifungals,^9^ with a well-established efficacy and safety profile.^8–12^ The primary mechanism of its antifungal action is the inhibition of ergosterol biosynthesis, by acting on the oxysterol-binding protein (OSBP).^13–16^ Since ergosterol is closely related to cholesterol, it has been suggested that also in mammalian cells, itraconazole acts as a cholesterol trafficking inhibitor resulting in disruption of the cholesterol-enriched membranes and thus inhibition of virus replication.^6,8,17^ However, a role in interferon priming has also been suggested as a contributing factor to the antiviral activity.^18^

In an attempt to identify therapeutic options for treating COVID-19 patients, the *in vitro* antiviral activities of itraconazole, and its metabolite 17-OH itraconazole were investigated in Caco-2 and VeroE6-eGFP cells infected with SARS-CoV-2, isolated from COVID-19 patients.

## Methods

### Cell culture

Caco-2 cells (human colon carcinoma cell line; obtained from the Deutsche Sammlung von Mikroorganismen und Zellkulturen, Braunschweig, Germany) were cultured in Minimal Essential Medium (MEM) supplemented with 10% fetal bovine serum (FBS) with penicillin (100 IU/mL) and streptomycin (100 μg/mL) at 37°C in a 5% CO_2_ atmosphere. VeroE6-eGFP (African green monkey kidney cell line; provided by Dr. K. Andries J&JPRD; Beerse, Belgium) were cultured in Dulbecco’s Modified Eagle Medium (DMEM) supplemented with 10% FBS, 0.075% sodium bicarbonate, penicillin and streptomycin (100 μg/mL) at 37°C in a 5% CO_2_ atmosphere. All cell culture reagents were obtained from Sigma-Aldrich (Hamburg, Germany).

### SARS-CoV-2 Preparation

SARS-CoV-2-FFM1 (strain hCoV-19/Germany/FrankfurtFFM1/2020) was isolated from a German human-case sample and cultured in Caco-2 cells, as previously described.^1,19^ SARS-CoV-2-Germany stocks were passaged twice in Caco-2 cells prior to storage (–80°C). SARS-CoV-2-Belgium (strain BetaCov/Belgium/GHB-03021/2020) was recovered from a nasopharyngeal swab taken from a patient returning from Wuhan, China. SARS-CoV-2 Belgium stocks were passaged six times in VeroE6-eGFP cells prior to storage (−80°C).

### Assessment of Antiviral Activity

Itraconazole and its metabolite, 17-OH itraconazole, and remdesivir were synthesized at Johnson and Johnson. GS-441524, the parent nucleoside of remdesivir, used in studies at KU Leuven was obtained from MedChemExpress (NJ, USA). Antiviral activity was assessed by inhibition of virus-induced cytopathogenic effect (CPE) as described previously.^20^ In brief, confluent layers of Caco-2 cells cultured for 72 hours on 96 multi-well plates (50,000 cells/well) were challenged with SARS-CoV-2 FFM1 at a multiplicity of infection of 0.01. The virus was added together with the compound under investigation and incubated in MEM supplemented with 1% FBS. Itraconazole diluted in MEM without FBS in 4-fold dilutions was added to a concentration range of 0.01 μM to 50 μM; 17-OH itraconazole diluted in MEM without FBS was added in 4-fold dilutions to a concentration range of 0.02 μM to 100 μM and remdesivir diluted in MEM without FBS in 4-fold dilutions was added to a concentration range of 0.02 μM to 100 μM. Cells were incubated for 48 hours; the CPE was then visually scored by two independent laboratory technicians. In addition, CPE was also assessed using a 3-(4,5-dimethylthiazol-2-yl)-2,5-diphenyltetrazolium bromide (MTT) assay, performed according to the manufacturer’s instructions. Optical densities were measured at 560/620 nm in a Multiskan Reader (MCC/340 Labsystems). Two series of three independent experiments (or one series of three experiments for 17OH-itraconazole), each containing three replicates, were performed. Data were analyzed by four-parameter curve-fitting from a dose-response curve using GraphPad Prism (version 7.00) to calculate the EC_50_ (concentration of the compound that inhibited 50% of the infection) based on visual CPE scoring or the MTT assay.

### Assessment of Cell Viability

Cell viability in Caco-2 cells was measured following administration of each of the compounds or metabolites under investigation over the range of concentrations in the absence of virus using the Rotitest Vital (Roth, Karlsruhe, Germany) test according to manufacturer’s instructions, as previously described.^21^ All assays were performed three times independently in triplicate, this was performed twice (once for 17OH-itraconazole). Data were analyzed by four-parameter curve-fitting from a dose-response curve using GraphPad Prism (version 7.00) to calculate the CC_50_ (cytotoxic concentration of the compound that reduced cell viability to 50%).

### Viral RNA Yield Reduction Assay

Antiviral activity was assessed by inhibition of viral yield in Caco-2 cells. Confluent layers of Caco-2 cells cultured for 72 hours on 96 multi-well plates (50,000 cells/well) were challenged with SARS-CoV-2-FFM1 at a multiplicity of infection of 0.01. The virus was added together with the compound under investigation and incubated in MEM supplemented with 1% FBS. Itraconazole diluted in MEM without FBS was added in 4-fold dilutions to a concentration range of 0.01 μM to 50 μM; 17-OH itraconazole diluted in MEM without FBS was added in 4-fold dilutions to a concentration range of 0.02 μM to 100 μM and remdesivir diluted in MEM without FBS was added in 4-fold dilutions to a concentration range of 0.02 μM to 100 μM. Two series of three independent experiments, each containing two replicates, were performed.

SARS-CoV-2 RNA from cell culture supernatant samples was isolated using AVL buffer and the QIAamp Viral RNA Kit (Qiagen) according to the manufacturer’s instructions. Absorbance-based quantification of the RNA yield was performed using the Genesys 10S UV-Vis Spectrophotometer (Thermo Scientific). RNA was subjected to OneStep qRT-PCR analysis using the Luna Universal One-Step RT-qPCR Kit (New England Biolabs) and a CFX96 Real-Time System, C1000 Touch Thermal Cycler (BioRad). Primers were adapted from the WHO protocol targeting the open reading frame for RNA-dependent RNA polymerase (RdRp): RdRP_SARSr-F2 (GTG ARA TGG TCA TGT GTG GCG G) and RdRP_SARSr-R1 (CAR ATG TTA AAS ACA CTA TTA GCA TA) using 0.4 μM per reaction. Standard curves were created using plasmid DNA (pEX-A128-RdRP) harboring the corresponding amplicon regions for RdRP target sequence according to GenBank Accession number NC_045512.

Antiviral activity was also assessed by reduction of viral yield in VeroE6-eGFP cells. VeroE6-eGFP cells were cultured for 24 hours on 96 multi-well plates (10,000 cells/well) in the absence or presence of compound and infected with SARS-CoV-2-Belgium at a multiplicity of infection of 1 Tissue Culture Infectious Dose (TCID_50_)/cell, in 200 μL assay medium (DMEM supplemented with 2% FBS and 0.075% sodium bicarbonate). After 2 hours at 37°C the cells were washed once and further cultured in 200 μL assay medium containing the same compound concentrations (37°C, 5% CO_2_) for another 48 hours. The viral RNA (vRNA) in the culture supernatant (SN) was extracted using the NucleoSpin kit (Macherey-Nagel), according to the manufacturer’s instructions and quantified by real-time quantitative polymerase chain reaction (RT-qPCR) performed on a LightCycler96 platform (Roche) using the iTaq Universal Probes One-Step RT-qPCR kit (BioRad) with primers and probes specific for SARS-CoV-2. A standard curve was created from serial dilutions of a virus stock with known titer and used to correlate the cycle threshold (Ct) values of the experimental samples with absolute virus quantities. One independent experiment, containing two replicates (for itraconazole) or three replicates (17OH-Itraconazole), were performed.

## Results

### *In Vitro* Antiviral Activity in Caco-2 cells

Independent experiments with triplicate measurements were performed with itraconazole (n=6), 17-OH itraconazole (n=3) and remdesivir (n=6). In Caco-2 cells, itraconazole resulted in a dose-dependent inhibition of SARS-CoV-2-FFM1 measured by visual scoring of inhibition of CPE with an EC_50_ = 1.5 μM (Figure 1). Similar findings were obtained using the MTT assay (EC_50_ = 2.3 μM). The 17-OH itraconazole metabolite reduced SARS-CoV-2 induced CPE with an EC_50 visual_ of 1.2 μM and EC_50 MTT_ = 3.6 μM. The positive control, remdesivir, resulted in EC_50_ values of 0.3 and 0.4 μM in the CPE and MTT assays in Caco-2 cells, respectively.

**Figure 1.**
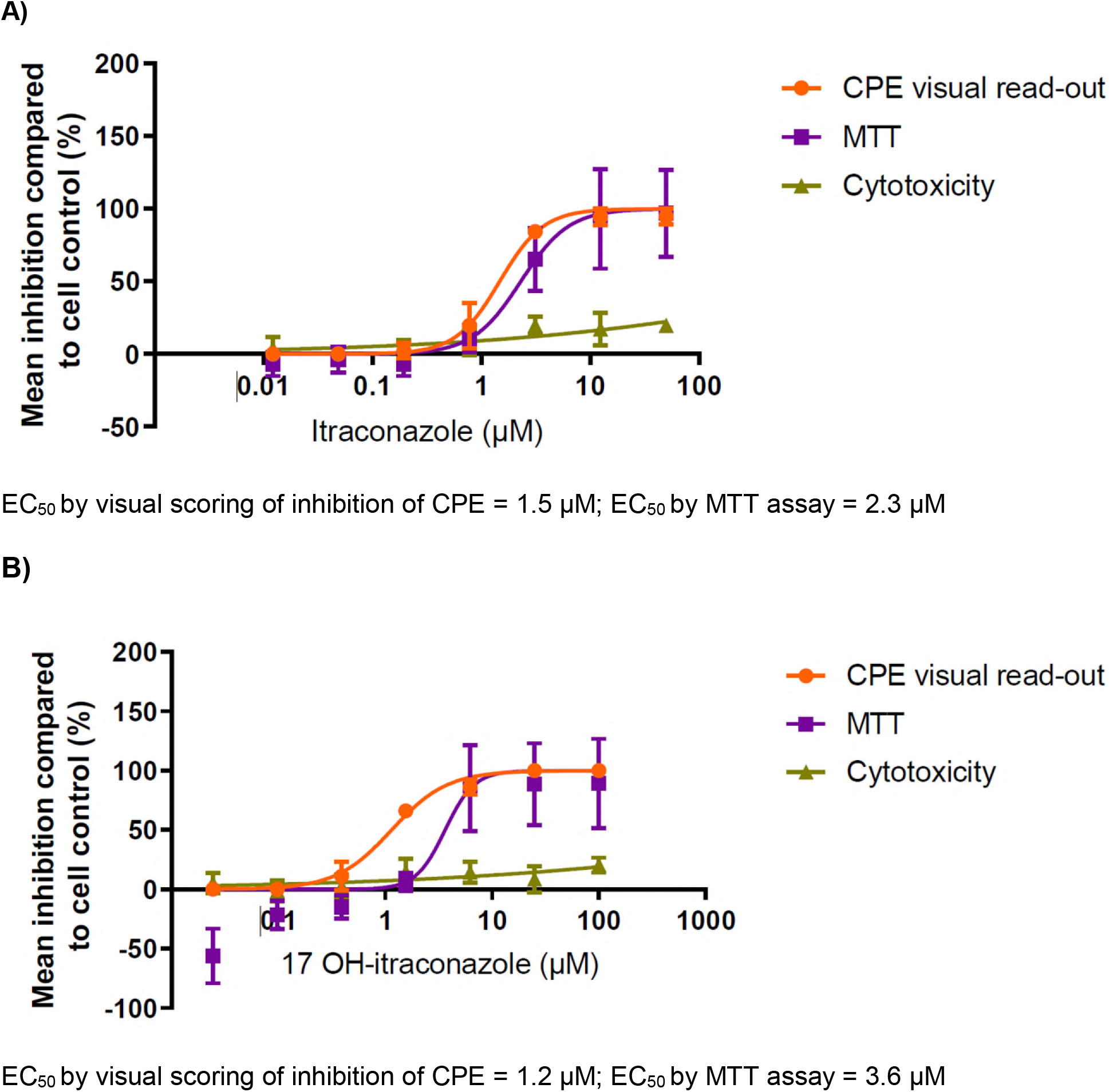
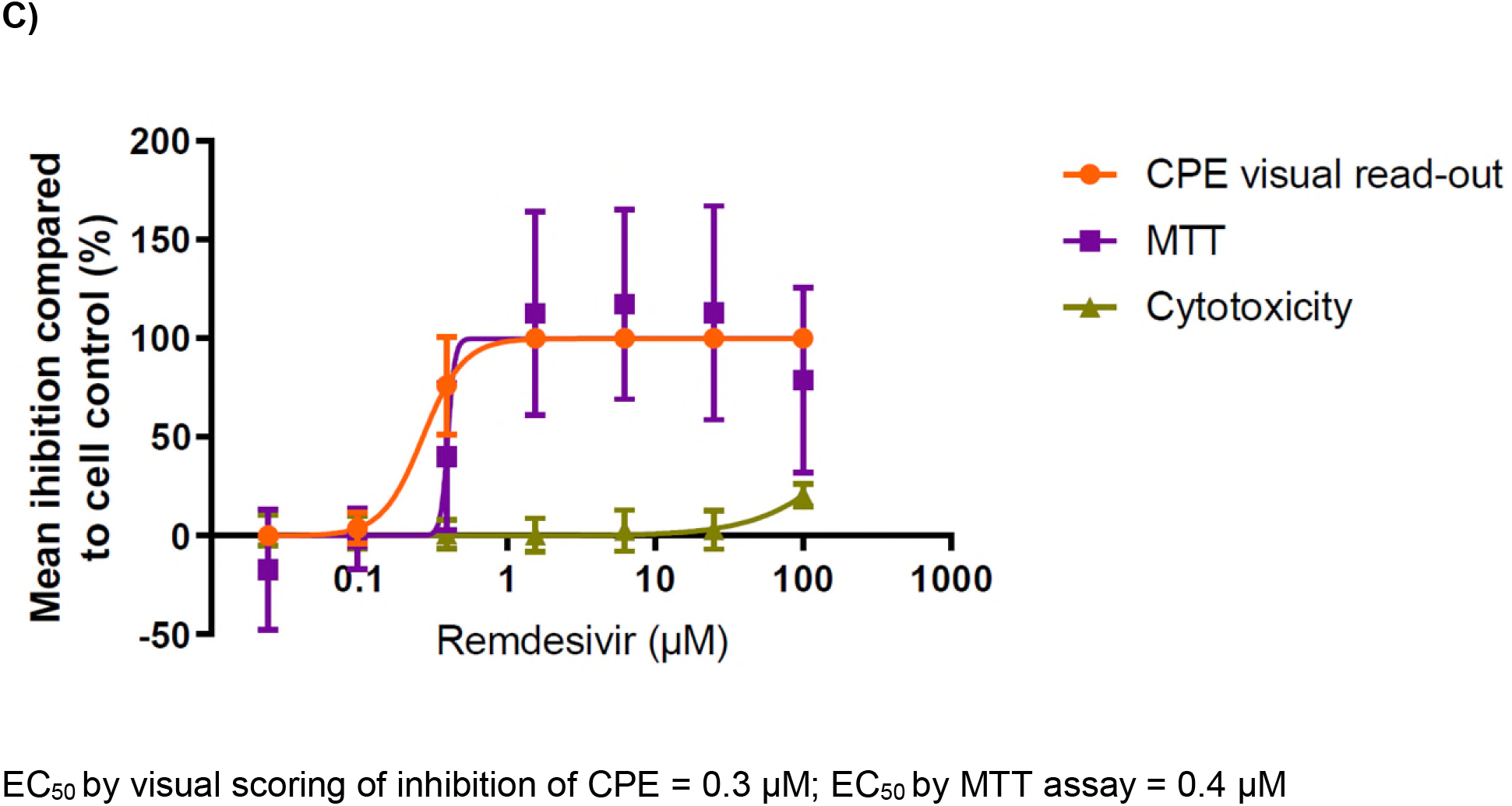
Effect of either itraconazole, 17-OH itraconazole or remdesivir on SARS-CoV-2 replication in CPE assays and on viability of Caco-2 cells. Mean percent inhibition for each readout across two series of three independent experiments (itraconazole, remdesivir) or three independent experiments (17-OH itraconazole) with triplicate measurements are plotted. The error bars represent the standard deviation. Orange represents CPE visual read-out; purple represents MTT assay; and green represents cytotoxicity.

Minimal cytotoxicity was seen in Caco-2 cells with both itraconazole (CC_50_ >50 μM) and 17-OH itraconazole (CC_50_ >100 μM) (Figure 1). For remdesivir CC_50_ values >100 μM were observed in Caco-2 cells.

### *In Vitro* Viral RNA Yield Reduction

The *in vitro* effect of itraconazole and remdesivir (Caco-2 cells) orGS-441522 (VeroE6-eGFP cells) on SARS-CoV-2 RNA yield reduction was assessed in Caco-2 cells and VeroE6-eGFP cells. A concentration of 6.25 μM and 3.1 μM of itraconazole and its main metabolite 17-OH itraconazole, respectively, resulted in an approximately 2-log_10_ reduction in viral RNA levels (as a measure of the number of virus particles in the culture SN) in Caco-2 cells. This reduction was observed at all higher doses, in the absence of toxicity. However, remdesivir proved more potent (a reduction of ~3 log_10_ at 6.25 μM) (Figure 2).

**Fig 2.**
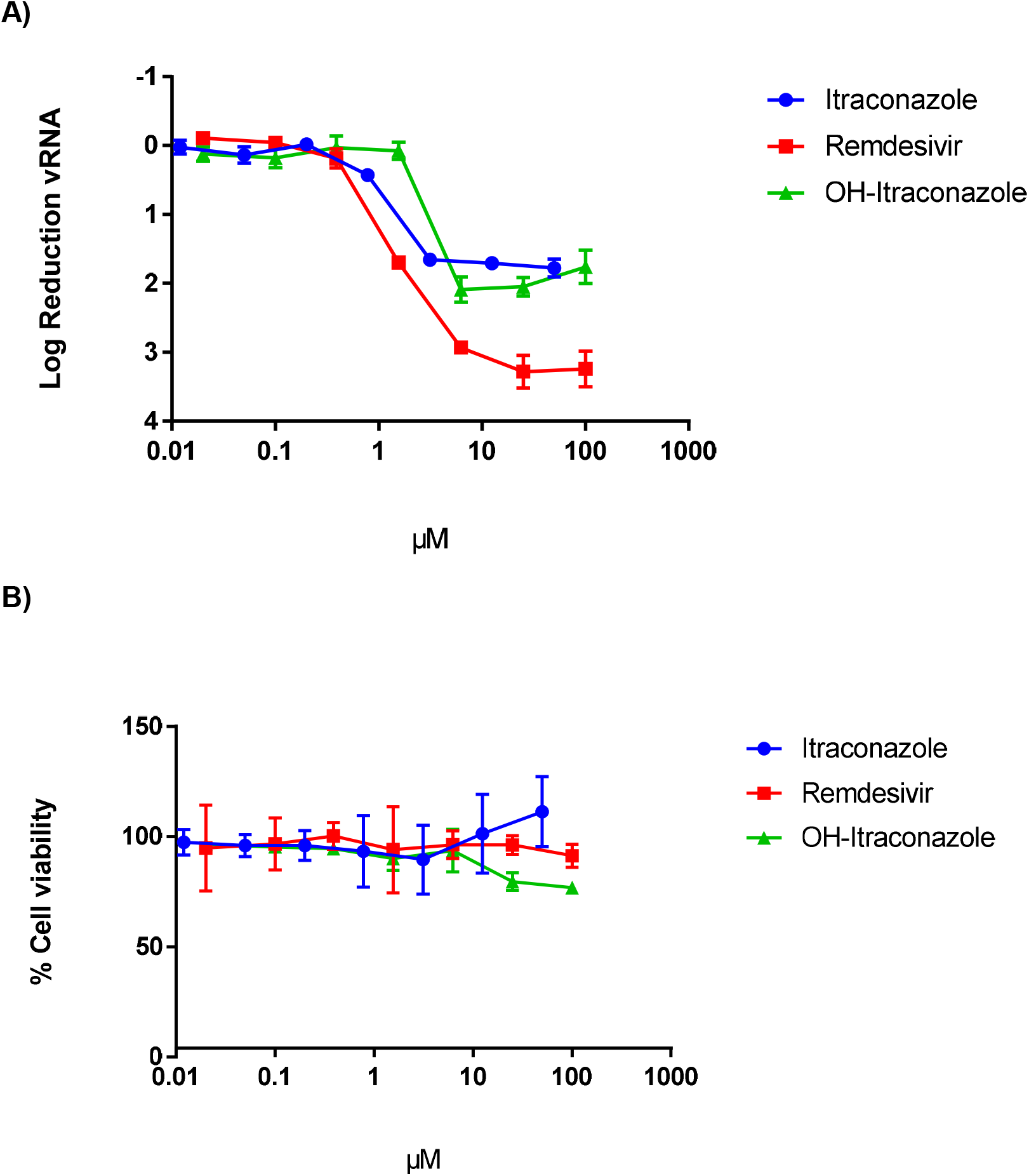
Effect of itraconazole, 17-OH itraconazole or remdesivir on SARS-CoV-2 vRNA yield and viability in Caco-2 cells. A) Mean differences in viral RNA in the supernatant between untreated cultures and treated cultures at 48 hours post-infection with SARS-CoV-2-FFM1 of three independent experiments each containing two replicates is shown. Error bars represent the standard deviation. B) Mean viability of the cells, based on three independent experiments each containing three replicates is shown. Error bars represent the standard deviation.

In VeroE6-eGFP cells, a concentration of 1 μM itraconazole resulted in a ~1-log_10_ reduction in viral RNA levels (Figure 3). This reduction was also observed at higher concentrations. As in Caco-2 cells, GS-441524 proved more potent in reducing viral RNA than itraconazole. At a concentration of 3.7 μM, GS-441524 reduced viral RNA load by >4 log_10_ to undetectable levels. At concentrations of 10 μM and higher, 17-OH itraconazole resulted in limited activity, but VeroE6-eGFP cell viability was markedly affected (Figure 3).

**Fig 3.**
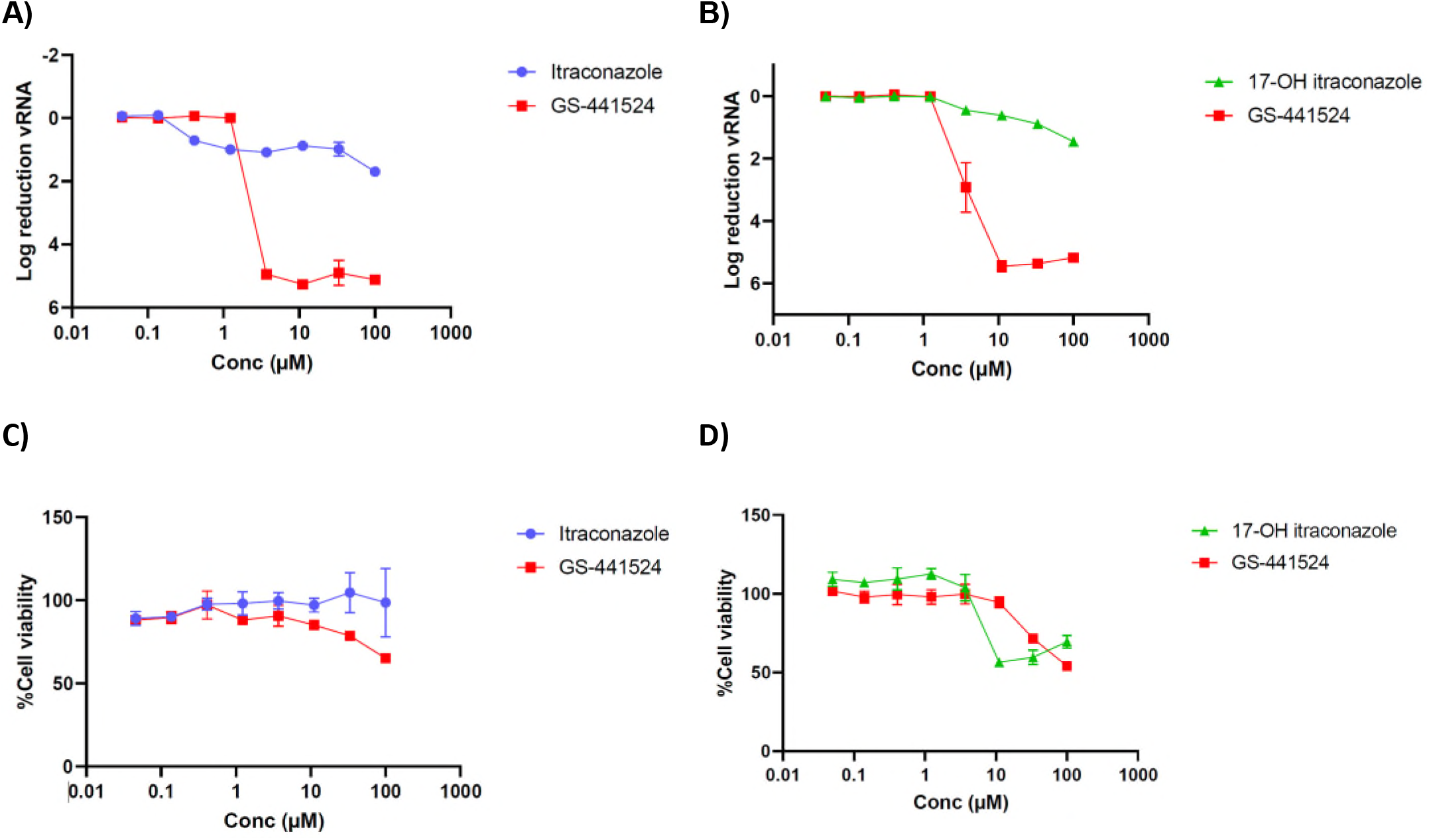
Effect of itraconazole, 17-OH itraconazole or GS-441524 on SARS-CoV-2 vRNA yield and viability in VeroE6-eGFP cells. A) and B) Mean differences in viral RNA in the supernatant between untreated cultures and treated cultures at 48 hours post-infection with SARS-CoV-2-Belgium of one independent experiment containing 2 replicates (A) or 3 replicates (B) is shown. Error bars represent the standard deviation. C) and D) Mean viability of the cells, based on MTT readout of uninfected cells. Error bars represent the standard deviation.

## Discussion

While a number of vaccines are in development, for the moment there are no antivirals for the prevention of COVID-19 and only a limited number of antivirals for the treatment of COVID-19.^22^ Given the great unmet need to identify potential treatments for COVID-19, and the fact that a *de novo* drug development project will take many years to bring a drug to patients, efforts have been expedited into the *in vitro* screening of compounds already on the market, or in late-stage development. In such an effort, itraconazole was identified, with single digit micromolar activity in Caco-2 cells. Itraconazole was first approved (in the US) in 1992 as an oral treatment for a number of fungal infections in immunocompromised and non-immunocompromised patients, with a well-established safety profile.^11^

These experiments demonstrated that itraconazole inhibits *in vitro* SARS-CoV-2 replication in Caco-2 cells. The EC_50_ values for inhibition of virus-induced CPE formation is in the low micromolar range and in that respect, is only around 5-fold higher than that of remdesivir. To obtain further evidence for the antiviral potency of the drug(s), the effect on viral RNA yield in culture was assessed in two cell lines. Itraconazole results in a marked inhibition of virus yield; however, itraconazole was less potent than remdesivir. Our findings are in line with recent data showing that itraconazole inhibited SARS-CoV-2 replication in a dose-dependent manner in both Calu-3 and Vero E6 cell lines (with EC_50_s of 0.43 μM and 0.39 μM respectively).^23^ In addition, *in vitro* studies have also demonstrated activity of itraconazole against murine coronavirus (recombinant murine coronavirus, mouse hepatitis virus, EC_50_ = 7.9 μM)^5^ and type I (but not type II) feline coronavirus (EC_50_ across three strains tested = 0.146–0.597 μM).^6^ Itraconazole has documented activity against a number of enteroviruses, including rhinovirus.^8,15^ Further, at a dose of 5.7 mg/kg, itraconazole resulted in improved mortality and a lower viral load in mice that had been infected with human influenza (strain PR8M) compared with the control group.^7^

As an antifungal agent, itraconazole inhibits fungal ergosterol biosynthesis, through inhibition of a cytochrome P450 enzyme, lanosterol-14-alpha-demethylase.^13–16^ It is also thought to interfere with cholesterol homeostasis and *de novo* synthesis in the host’s cells through inhibition of this same enzyme, thereby influencing the virus/host cell interaction.^17,24–26^ The antiviral effect of itraconazole may be, at least in part, based on such a mechanism. Itraconazole impairs cholesterol trafficking and blocks late endosomal/lysosomal export of cholesterol to the plasma membrane through NPC1 leading to cholesterol accumulation in sub-cellular compartments.^8,16,24–27^ In turn, elevated cholesterol levels in the endosomal membrane impede fusion of the viral lipid envelope and prevent viral genome transfer into the host cell cytosol.^7,28^ It has been postulated that such changes in cellular cholesterol are linked with the host’s immune response.^29^ Disruption of cholesterol biosynthesis has been shown to upregulate type I interferons and thereby accelerate the virus-induced interferon-mediated host cell response.^7,29^ It has also been suggested that itraconazole may have antiviral effects through proteins other than NPC intracellular cholesterol transporter 1 (NPC1).^6^ In enteroviruses, itraconazole has been shown to inhibit viral RNA replication by oxysterol-binding protein (OSBP), which is responsible for trafficking of cholesterol and phosphatidylinositol-4-phosphate between membranes, thus also affecting the membranes of the replication complex.^15,16^ Since coronaviruses result in the formation of intracellular viral replicative organelles within the endoplasmic reticulum that drive the viral replication cycle,^30–32^ it is possible that inhibition of OSBP may cause detrimental changes to replicative organelle membranes.^15,16,32,33^ Although it remains to be confirmed how itraconazole may inhibit the replication cycle of coronaviruses, it is conceivable that it acts through multiple mechanisms.^7,17^

As a lipophilic compound, itraconazole has high bioavailability and extensive distribution throughout the lung, kidney, epidermis and brain.^9^ Itraconazole is predominantly metabolized by the cytochrome P450 3A4 isoenzyme system, forming many metabolites.^34,35^ The main metabolite produced is 17-OH itraconazole, which also has considerable antifungal activity.^34^ Following administration of itraconazole 200mg bid, itraconazole and its 17-OH hydroxy-metabolite have been shown to reach mean maximal plasma concentrations of 2282 and 3488 ng/mL (~ 3.2 μM and ~4.8 μM) and a C_trough_ of 1855 and 3349 ng/ml (~2.6 μM and ~4.7 μM), respectively, which is close to the EC_50_ for inhibition of virus-induced CPE. Both the parent drug and metabolite have long half-lives (64 and 56 hours, respectively).^11^

In conclusion, itraconazole and its metabolite, 17-OH itraconazole show *in vitro* activity against SARS-CoV-2. Based on these *in vitro* findings, a clinical study in hospitalized COVID-19 patients was initiated by UZ Leuven in March 2020.^36^ However, no efficacy of the drug was observed in this particular study population; other studies in non-hospitalized patients were not conducted.

## Acknowledgments

This project has received funding from the European Union’s Horizon 2020 research and innovation program under grant agreement no.: 101003627. Part of this research work was performed using the ‘Caps-It’ research infrastructure (project ZW13-02) that was financially supported by the Hercules Foundation and Rega Foundation, KU, Leuven. This work has been funded in part with Federal funds from the Office of the Assistant Secretary for Preparedness and Response, Biomedical Advanced Research and Development Authority, under OTA No. HHSO100201800012C. Medical writing supporting for the development of this manuscript was provided by Patrick Hoggard of Zoetic Science, an Ashfield company, part of UDG Healthcare plc; this support was funded by Janssen Pharmaceuticals. We thank Lena Stegmann, Winston Chiu, Piet Maes, Robbert Boudewijns, Nguyen Dan Thuc Do, Xin Zhang, Jana Van Dycke, Rana Abdelnabi, Laura Thijs, Peggy Geluykens, Doortje Borrenberghs, and Christel Van den Eynde for technical support for antiviral assays.

## Disclaimers

Sandra De Meyer, Ellen Van Damme, Christophe Buyck and Marnix Van Loock are employees and may be stock owners of Johnson & Johnson. Sandra Ciesek received research funding from Janssen for this research. Steven De Jonghe, Dirk Jochmans, Pieter Leyssen and Johan Neyts are employees from KU Leuven and received funding from Janssen for this research.

The data sharing policy of Janssen Pharmaceutical Companies of Johnson & Johnson is available at https://www.janssen.com/clinical-trials/transparency. As noted on this site, requests for access to the study data can be submitted through Yale Open Data Access (YODA) Project site at http://yoda.yale.edu.

